# Unstable inheritance of 45S rRNA genes in *Arabidopsis thaliana*

**DOI:** 10.1101/088088

**Authors:** Fernando A. Rabanal, Viktoria Nizhynska, Terezie Mandáková, Polina Yu. Novikova, Martin A. Lysak, Richard Mott, Magnus Nordborg

## Abstract

The considerable genome size variation in *Arabidopsis thaliana* has been shown largely to be due to copy number variation (CNV) in 45S ribosomal RNA (rRNA) genes. Surprisingly, attempts to map this variation by means of genome-wide association studies (GWAS) failed to identify either of the two likely sources, namely the nucleolar organizer regions (NORs). Instead, GWAS implicated a *trans*-acting locus, as if rRNA CNV was a phenotype rather than a genotype. To explain these results, we investigated the inheritance and stability of rRNA gene copy number using the variety of genetic resources available in *A. thaliana* — F2 crosses, recombinant inbred lines, the multiparent advanced generation inter-cross population, and mutation accumulation lines. Our results clearly show that rRNA gene CNV can be mapped to the NORs themselves, with both loci contributing equally to the variation. However, NOR size is unstably inherited, and dramatic copy number changes are visible already within tens of generations, which explains why it is not possible to map the NORs using GWAS. We did not find any evidence of *trans*-acting loci in crosses, which is also expected since changes due to such loci would take very many generations to manifest themselves. rRNA gene copy number is thus an interesting example of “missing heritability” — a trait that is heritable in pedigrees, but not in the general population.

## Introduction

In eukaryotic genomes, 45S rRNA genes are arranged in clusters termed nucleolus organizer regions (NORs) (Long and Dawid 1980). After transcription by RNA polymerase I, the primary transcript is processed into 18S, 5.8S and 25S rRNAs that, together with the 5S rRNA (encoded by a separate multi-copy gene), constitute the catalytic core of ribosomes (Chambon 1975; Long and Dawid 1980). In *A. thaliana*, each 45S ribosomal RNA (rRNA) gene is over 10 kb long, and the genome contains hundreds of tandemly arrayed gene copies at the top of chromosomes 2 (NOR2) and 4 (NOR4) (Copenhaver *et al.* 1995; Copenhaver and Pikaard 1996a). Natural inbred lines (accessions) vary by well over 10% in genome size (Schmuths *et al.* 2004; Long *et al.* 2013), largely due to differences in 45S rRNA gene copy number (Davison *et al.* 2007; Long *et al.* 2013). However, besides pulsed-field electrophoresis studies in the accession Landsberg indicating that both NORs are similar in size, each spanning approximately 3.5-4.0 Mb (Copenhaver and Pikaard 1996b), nothing is known about the specific contribution of each locus to the overall copy number variation (CNV) in 45S rRNA genes.

We previously carried out a genome-wide association study (GWAS) to investigate the genetics of both the variation in genome size and 45S rRNA gene CNV in a population of *A. thaliana* lines from Sweden. We expected to find significant associations in *cis* — due to strong linkage disequilibrium between NOR haplotypes and closely linked single nucleotide polymorphisms (SNPs). Surprisingly, the scans identified neither of the two NORs. Instead, the analyses found an association in *trans* on chromosome 1, as if rRNA gene copy number were a phenotype rather than a genotype (Long *et al.* 2013).

Alternatively, repeat number may change too rapidly to be mapped using GWAS, but may still be inherited stably enough to be mapped in crosses (Long *et al.* 2013). Consistent with this, quantitative trait locus (QTL) analyses aimed at understanding the genetics behind NOR methylation in *A. thaliana* have suggested that CNV at the NORs themselves accounts for some of the methylation variation (Riddle and Richards 2002a, 2005). Indeed, rapid changes in 45S rRNA gene copy number have been detected for several species. Examples range from a ∼2-fold variation in copy number after 400 generations in fruit fly lines and nematodes (Averbeck and Eickbush 2005; Bik *et al.* 2013) or a similar 2.5-fold variation after only 70 generations in maize lines (Phillips 1978), to differences greater than 4-fold after 90 generations in water flea lines (McTaggart *et al.* 2007) or even greater than 2-fold changes across siblings in humans (Gibbons *et al.* 2015) or 7-fold changes among individual siblings of a self-pollinated faba bean parent (Rogers and Bendich 1987). In light of the various degrees of instability in rRNA gene copy number displayed by higher plants (Walbot and Cullis 1985), it is relevant to investigate how rapidly the number of rRNA genes changes in *A. thaliana*.

Our aim in this study was threefold: first, to test if the *trans* association detected by GWAS (Long *et al.* 2013) has an effect in a segregating F2 population; second, to confirm that CNV in rRNA genes can be mapped to the NORs themselves in crosses; third, to investigate how copy number in rRNA genes of *A. thaliana* changes on a generational time scale.

## Results

### 45S rRNA gene CNV can be mapped to specific NORs in F2s

To better understand the genetics of 45S rRNA gene CNV, we generated an F2 population from a cross between a large copy number accession from northern Sweden — TRÄ-01 (6244), with ∼2,500 units — and a small copy number accession from southern Sweden — Ale-Stenar-64-24 (1002), with ∼500 units. We used next generation sequencing (NGS) to phenotype (we estimated the copy number of the 18S rRNA gene, which is strongly correlated with the copy number of the full gene) and genotype the population simultaneously (Figure 1A; see Methods). In sharp contrast to GWAS, linkage mapping identified the distal end region at the top of chromosome 2 as the sole source of variation in rRNA gene copy number in this population (Figure 1B). The *trans*-association identified by GWAS in chromosome 1 (Long *et al.* 2013) was not captured by this analysis, despite the fact that the alleles responsible for the presumed association segregate in the parental accessions.

**Figure 1.**
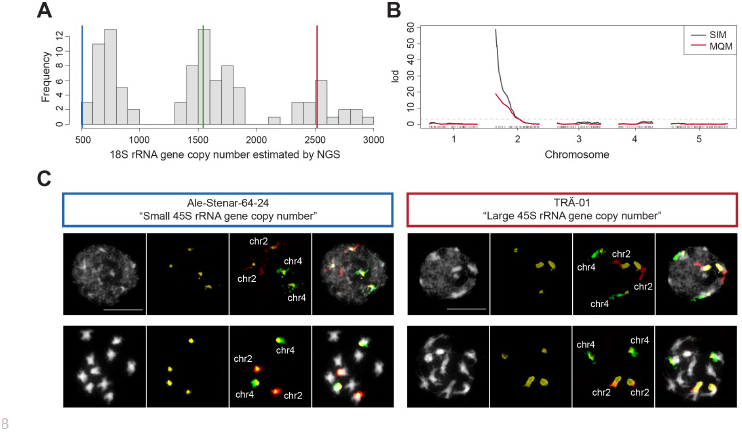
rRNA gene copy number variation in an F2 population is driven by NOR2. (A) The distribution of 18S rRNA gene copy number estimated by NGS in an F2 population of 93 individuals derived from the cross Ale-Stenar-64-24 (1002) × TRÄ-01 (6244). Blue, green and red vertical lines represent phenotypic values of accession Ale-Stenar-64-24, an F1 individual and accession TRÄ-01, respectively. (B) QTL mapping of 18S rRNA gene copy number in the same F2 population. Black and red lines indicate simple interval mapping (SIM) and multiple-QTL mapping (MQM) models, respectively (Broman *et al.* 2003; Arends *et al.* 2010). (C) FISH results for the parental lines Ale-Stenar-64-24 and TRÄ-01. Images in black and white show DAPI-stained nuclei (upper panels) and mitotic chromosomes (lower panels). Probes hybridizing the 45S rRNA gene cluster, chromosomes 2 and 4 are highlighted in yellow, red and green, respectively. Bar = 10 μm.

To corroborate that NOR2 is indeed responsible for the difference in rRNA gene copy number, we performed fluorescence in situ hybridization (FISH) in both parental accessions. The results showed that NOR2 and NOR4 in the southern accession Ale-Stenar-64-24 are of similar size to each other — NOR4 is on average 1.49x larger (106.5/71.17 pixels; n=29) than NOR2 in mitotic chromosomes — while in the northern accession TRÄ-01, NOR2 is 2.39x larger than NOR4 (299.64/125.27 pixels; n=26) (Figure 1C).

Mapping in two further F2 populations showed that it is not always NOR2 varying in size. CNV mapped to NOR2 in the cross Ull1-1 (8426) × TDr-7 (6193) (Figure S1, A and B), but to NOR4 in the cross T460 (6106) × Omn-5 (6071) (Figure S1, C and D). In neither population was there evidence of any *trans*-acting loci. Taken together, these results show that both NORs vary in size, and that this size is stable enough to be readily traced over two generations. Given this stability, it is not unexpected that we saw no evidence of *trans*-acting loci, because such loci would by necessity modify the copy number.

### Size heterogeneity of rRNA gene loci in a worldwide population

Two of our F2 populations identified NOR2 as the major source of CNV; one identified NOR4. To improve our understanding of CNV in the general population, beyond a few biparental crosses, we employed the multi-parent advanced-generation inter-cross (MAGIC) population that is derived from intercrossing 19 world-wide accession (Kover *et al.* 2009). Mapping of 18S rRNA gene copy number in 393 individuals of the MAGIC population revealed that both NORs contribute to the variation to a similar extent (Figure 2A), with the contribution varying greatly among founder lines (Figure 2B). For example, on average, MAGIC lines carrying NOR2 from accessions Bur-0 (7058) and Zu-0 (7417) have fewer copies than do lines that carry NOR4 from these lines instead, because — as confirmed by FISH — founder accessions Bur-0 (Figure 2C) and Zu-0 (Figure 2D) have larger NOR4 than NOR2. Remarkably, we were unable to detect any fluorescence corresponding to 45S rRNA genes in chromosome 2 of Bur-0, suggesting that NOR2 is almost absent in this line (Figure 2C).

**Figure 2.**
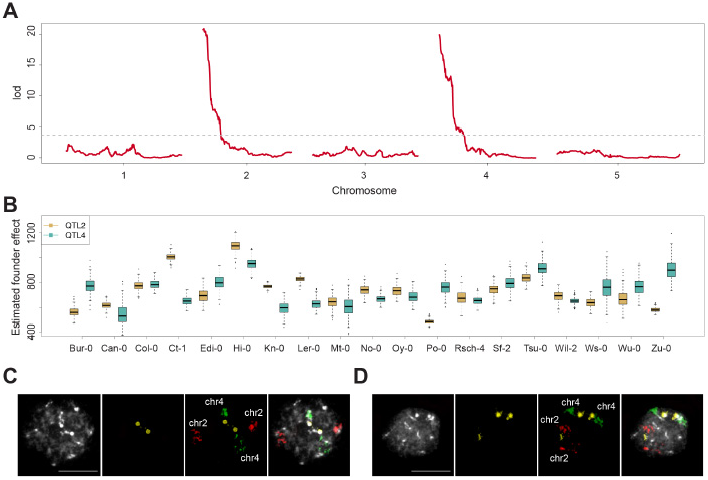
Mapping in the MAGIC lines reveals both NORs in *A. thaliana* contribute to the variation in rRNA gene copy number. (A) QTL mapping of 18S rRNA gene copy number variation in 393 individuals of the MAGIC population estimated by NGS. (B) Estimated founder accession effect by multiple imputation using R/happy (Mott *et al.* 2000; Kover *et al.* 2009) at significant QTLs on both chromosomes 2 and 4. (C) FISH results for the founder line Bur-0. Images in black and white show DAPI-stained nuclei. Probes hybridizing the 45S rRNA gene cluster, chromosomes 2 and 4 are highlighted in yellow, red and green fluorescence, respectively. Bar = 10 μm. (D) FISH results for the founder line Zu-0 as described in (C).

### Unstable inheritance of rRNA gene copy number in a RIL population

While rRNA gene copy number appeared stable in F2 progeny (Figure 1 and Figure S1, A-D), we thought it might be possible to observe changes in recombinant inbred line (RIL) populations, which have typically undergone at least eight generations of inbreeding since the original cross. Mapping in a RIL population derived from a cross between Cvi-0 and Ler-0 (Alonso-Blanco *et al.* 1998) — two accessions that differ by as few as ∼100 rRNA gene copies (Riddle and Richards 2002a) (Figure S1E) — showed that rRNA gene CNV maps to NOR2 (Figure S1F). However, after splitting the estimates of rRNA gene copy number by parental origin for each NOR, aberrant values became apparent (Figure 3A). Most notably, CVL45 carried ∼200 rRNA gene copies less than other individuals with Cvi-only NORs, while CVL168 and CVL102 have ∼150 and ∼250 fewer copies, respectively, than other individuals carrying Ler-only NORs (Figure 3A).

**Figure 3.**
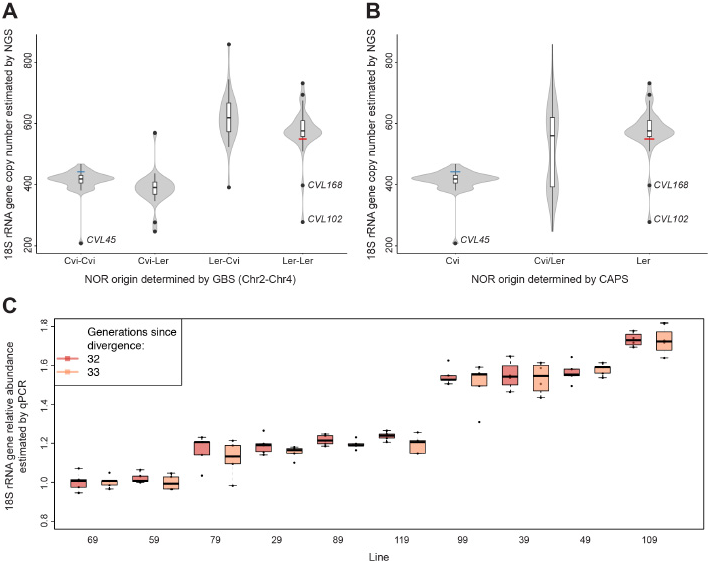
Instability of the rRNA gene repeats is manifested in a small number of generations. (A) 18S rRNA gene copy number in the Cvi-0 × Ler-0 RIL population estimated by NGS split by NOR parental identity as determined by genotyping by sequencing (GBS). (B) 18S rRNA gene copy number in the Cvi-0 × Ler-0 RIL population estimated by NGS split by NOR parental identity as determined by CAPS assay. (C) 18S rRNA gene copy number in the Mutation Accumulation lines estimated by qPCR in two consecutive generations (32 and 33).

To rule out that these drastic changes were due to interchromosomal exchange (recombination) between homologous NORs of different parental origin, we performed CAPS analysis that discriminates between rRNA genes of the parental accessions Cvi-0 and Ler-0 (Lewis *et al.* 2004) (Figure 3B). This analysis revealed that the low copy number phenotypes of CVL102 and CVL168 cannot be the product of recombination with Cvi-0 NORs, since no traces of Cvi-like NORs were identified. Similarly, CVL45 contains exclusively Cvi-0 NORs (Figure 3B). Copy number must thus have mutated in these lines, perhaps via unequal crossing-over. Indeed, our observations are consistent with numerous studies suggesting that unequal crossing over is the prevalent mechanism in the evolution and dynamics of rRNA genes (Eickbush and Eickbush 2007); with sister chromatid exchange being more frequent than exchange between homologs in budding yeast (Petes 1980; Szostak and Wu 1980), fruit flies (Williams *et al.* 1989; Schlötterer and Tautz 1994) and humans (Seperack et al. 1988). Worth noticing is that the distribution of rRNA gene copy number in this RIL population, which has undergone at least 9 generations of inbreeding, shows an apparent lack of F1-like phenotypes (Figure S1E), further supporting the notion that NORs in homologous chromosomes do not readily recombine in *A. thaliana* (Copenhaver *et al.* 1995). This is in apparent contrast to humans, where presumably meiotic recombination accounts for the striking variability observed at single NORs in parent-child trios (Schmickel *et al.* 1985; Kuick *et al.* 1996; Stults *et al.* 2008).

Changes in rRNA gene copy number may be associated with changes in heterochromatin formation (Paredes and Maggert 2009). Relative to Ler-0, Cvi-0 has reduced chromatin compaction, and QTL mapping (using the same RIL population used here), pointed to *PHYTOCHROME-B* (PHYB) and *HISTONE DEACETYLASE 6* (HDA6) as regulators of light-mediated chromatin compaction (Tessadori *et al.* 2009).

Furthermore, the decreased levels of DNA and histone H3K9 methylation at the NORs resembled those seen in the *hda6* mutant in the Col-0 background (Riddle and Richards 2002b; Earley *et al.* 2006, 2010; Tessadori *et al.* 2009). Although our mapping did not identify significant *trans*-acting QTL for rRNA gene CNV in this RIL population (Figure S1F), we tested the effect of NOR-of-origin as a function of the allele (Ler-0 or Cvi-0) inherited at either *PHYB* or *HDA6* directly (using a linear model). This analysis revealed no significant contribution of *PHYB* (Figure S2A), and only a marginally significant interaction for the role of HDA6 at NOR genotypes Ler-Cvi and Ler-Ler — p-value = 0.0298 and p-value = 0.0125, respectively (Figure S2B).

### Unstable inheritance of rRNA gene copy number in mutation accumulation lines

We next turned to mutation accumulation (MA) lines: independent descendants of the reference accession Col-0 that have been maintained by single-seed descent for over 30 generations in the absence of selection (Shaw *et al.* 2000). Note that since these are inbred lines, changes in copy number due to recombination between copy-number variants can definitely be ruled out. We quantified 18S rRNA gene copy number by qPCR for two consecutive generations in ten lines that have diverged for 31 generations (Ossowski *et al.* 2010; Schmitz *et al.* 2011; Becker *et al.* 2011) (Figure 3C). We considered a full linear mixed-effects model in which ‘line’ and ‘generation’ were added as fixed effects, while ‘replicates’ per line across generations were added as random effects. We used likelihood ratio tests to compare the full model and two reduced models: (1) omitting ‘line’ — the effect of 30 generations since divergence — or; (2) ‘generation’ — the effect of one subsequent propagation by single seed descent. While ‘line’ significantly affected rRNA copy number (χ^2^ (1)=298.19, p-value < 2.2e-16), ‘generation’ had a negligible impact (χ^2^ (2)=3.6, p-value = 0.057). In other words, the difference among independent MA lines accumulated in the 31 generations since divergence is much greater than the one manifested in only one generation — or the intrinsic error of our measurement. That these estimates are reliable is also evidenced by the good correlation between qPCR and NGS estimates for generation 31 (R-squared = 0.88, p-value = 6.105e-05; Figure S3) (Becker *et al.* 2011; Hagmann *et al.* 2015). There is thus clear evidence for instability of rRNA gene copy number over as few as 30 generations.

## Discussion

This study was motivated by our observation that rRNA gene copy number, the major determinant of genome size variation in *A. thaliana*, behaved very strangely in GWAS (Long et al. 2013). Specifically, although the variation was likely to be due to CNV at the NORs, we were not able to map them in *cis*. Instead, we mapped what appeared to be a *trans*-acting locus, which prompted us to consider rRNA gene CNV as a phenotype rather than a genotype, at least in part (Long *et al.* 2013). To help make sense of these findings, we decided to study the pattern of inheritance using F2s and inbred lines. As opposed to the case in humans (Schmickel *et al.* 1985; Kuick *et al.* 1996; Stults *et al.* 2008), we found that rRNA gene copy number clearly behaves like a genetic trait in pedigrees, with the trait mapping either to NOR2 or NOR4 depending on the parents (Figure 1, Figure S1 and Figure 2). However, we also found that the trait is unstably inherited: by amassing estimates of rRNA gene copy number from F2s, RILs and MA lines in sets of individuals sharing the same genotypes at both NORs, we were able to show that progressive copy number changes are evident already in tens of generations (Figure 4). Together, these two observations provide an explanation for why we were not able to map the NORs using GWAS: copy number is simply too unstable, and hence not heritable over the time scales relevant in GWAS. This is thus a *bona fide* case of “missing heritability” — a trait that is heritable in families, but cannot possibly be mapped using GWAS (Manolio *et al.* 2009).

**Figure 4.**
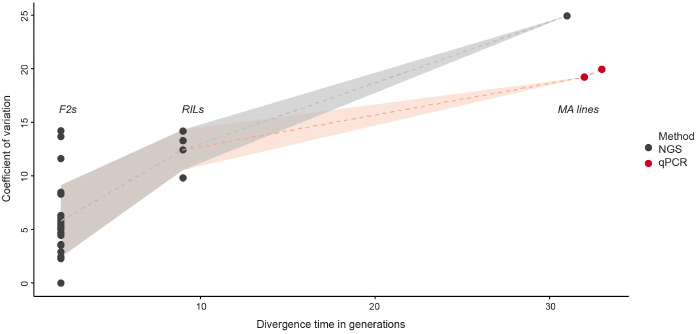
Coefficient of variation in rRNA gene copy number along generations since divergence. Coefficient of variation in rRNA gene copy number along generations since divergence for sets of individuals sharing the same genotypes at both NOR loci. For generations 2 and 9 data was collected from F2 and RIL populations, respectively; while for the latest generations data was collected from Mutation Accumulation lines. Black and red dots represent estimates by NGS and qPCR, respectively.

We did not find any evidence for *trans*-acting loci affecting rRNA gene copy number in any of the artificial mapping populations used in this study. Since we now know that the trait behaves like a genotype rather than a phenotype on this time scale, this is not surprising. It also does not imply that the reported association (Long *et al.* 2013) is a false positive, because a trans-acting locus that works by biasing the mutation process, predisposing carriers to acquire more or less copies, would not have any effect over a few generations. Such a locus may still affect genome size in local populations of *A. thaliana*, and be mappable using GWAS (this would thus be exactly the opposite of missing heritability — a phenotype that is only heritable on a population scale but cannot be observed in pedigrees). Resolving this through crosses may be difficult in a plant with relatively long life cycle.

Does all this variation have any biological relevance? It has been recently shown that in *A. thaliana* Col-0, only NOR4 derived rRNA genes are actively transcribed and associated with the nucleolus, while NOR2 is silent (Pontvianne *et al.* 2010, 2013; Chandrasekhara *et al.* 2016). However, our cytological analysis showed that, in TRÄ-01, NOR2 is the NOR associated with the nucleolus, indicating that in this accession NOR2 rRNA genes might be the active ones (Figure 1C, Table S1). Furthermore, using transcriptome analysis (F.A. Rabanal and M. Nordborg, *manuscript in preparation*, File S3), we identified great variation among accessions in which NOR is utilized, and demonstrated that a complex dominance hierarchy appears to exists among NOR haplotypes. Thus, not only do both clusters contribute to genome size variation (Figure 1, Figure S1 and Figure 2), they also contribute to rRNA expression in natural populations. Why this should be the case remains unclear.

In conclusion, we have shown that rRNA gene copy number is semi-conservatively inherited and starts to diverge over a time-scale of tens of generations. As a result, the trait is heritable in pedigrees, but cannot be mapped using GWAS. This resolves the seemingly paradoxical GWAS results for rRNA gene CNV in *A. thaliana*, and lays the ground for trying to understand whether any of the observed variation has functional importance, as suggested by its geographic distribution (Long *et al.* 2013).

## Materials and Methods

### DNA extraction and library preparation

We harvested leaves from ∼3 weeks old plants grown under long day conditions (16 hrs. light and 8 hrs. at 10°C). We extracted DNA in 96-well plates with the NucleoMag^®^ 96 Plant (Macherey-Nagel) kit according to the manufacturer's instructions.

We prepared libraries using a slightly modified version of the Illumina Genomic DNA Sample preparation protocol. Briefly, 100 to 200 ng of DNA were fragmented by sonication with Bioruptor (Diagenode). End-repair of sheared DNA fragments, A-tailing and adapter ligation were done with Spark DNA Sample Prep Kit (Enzymatics). NEXTflex-96™ DNA Barcodes (Bioo Scientific) were used to attach indexes to the sample insert during adapter ligation. Size selection, with median insert size around 400 bp, and library purification were performed with Agencourt AMPure XP Beads (Beckman Coulter). Paired-end (PE) DNA libraries were amplified by PCR for 10–12 cycles. After PCR enrichment, libraries were validated with Fragment Analyzer™ Automated CE System (Advanced Analytical) and pooled in equimolar concentration for 96X-multiplex. Libraries were sequenced on Illumina HiSeq™ 2000 Analyzers using manufacturer's standard cluster generation and sequencing protocols in 100 bp PE mode.

### Genotyping by sequencing

For each segregating F2 or RIL population analysed in this study (1002×6244, 6106×6071, 8426×6193, 6911×7213) we applied the following pipeline separately. We extracted both known indels and biallelic homozygous SNPs of the parental accessions from the 1001 Genomes Consortium (1001 Genomes Consortium 2016) with SelectVariants from Genome Analysis Toolkit (GATK; v3.5) (DePristo *et al.* 2011; Van der Auwera *et al.* 2013). We combined only segregating SNPs between parental accessions in a single variant call format (VCF) file with GATK/CombineVariants for later genotyping of individual samples (see below).

For each low-coverage sample we mapped PE reads to the *Arabidopsis thaliana* TAIR10 reference genome with BWA-MEM (v0.7.4) (Li and Durbin 2009; Li 2013). We used Samtools (v0.1.18) to convert file formats, sort and index bam files (Li *et al.* 2009), while to remove duplicated reads we used Markduplicates from Picard (v1.101) (http://broadinstitute.github.io/picard/). We performed local realignment around indels by providing to the GATK/RealignerTargetCreator function known indels from the parental accessions to generate the set of intervals required by the GATK/IndelRealigner function. We called SNPs at the segregating sites determined in the combined VCF of the parental accessions with GATK/UnifiedGenotyper in genotyping mode with parameters ‘-glm SNP -gt_mode GENOTYPE_GIVEN_ALLELES -stand_call_conf 0.0 – G none -out_mode EMIT_ALL_SITES’.

For the construction of individual genetic maps we binned marker SNPs in 100 kb windows using R software with help of the package R/xts (Ryan and Ulrich 2011; Team 2014). We discarded windows with either less than 100 segregating SNPs or less than 40 called SNPs. The former for considering them regions of low diversity between parental accessions, while the latter for considering them regions not well supported by reads. We assigned genotype ‘A’ or genotype ‘B’ to windows with more than 90% of SNP calls for the maternal or paternal accessions, respectively. We determined as genotype ‘H’ windows with either more than 25% heterozygous calls or where the absolute difference between maternal and paternal SNP calls were less than 30%.

### Estimating rRNA gene copy number through NGS

For each individual we mapped all reads separately to a single reference 45S rRNA gene (extracted from GenBank: CP002686.1 coordinates 14195483–14204860; File S1) and to the A. *thaliana* TAIR10 reference genome as described in the section ‘Genotyping by sequencing’. For our reference 45S rRNA gene (File S1), we based the annotations of the 18S, 5.8S and 25S subunits (coordinates 2195–4002, 4271–4434 and 4623–8009, respectively) in previous reports (Gruendler *et al.* 1989; Unfried *et al.* 1989; Unfried and Gruendler 1990; Cokus *et al.* 2008). We retrieved per-base read depth with the function Depthofcoverage from GATK (Van der Auwera *et al.* 2013) before and after removal of duplicated reads. Since the correlation between NGS and qPCR estimates of 45S rRNA gene copy number has been shown to be better before removal of duplicated reads (Long *et al.* 2013), we performed further quantitative analysis with NGS estimates accordingly.

Since estimates of the 18S and 25S subunits of the 45S rRNA gene are in good agreement (Davison *et al.* 2007), we estimated 45S rRNA gene copy number in F2s, RILs, MAGIC lines and MA lines through next generation sequencing (NGS) by dividing the average coverage along the 18S rRNA gene by the average coverage along the first 10 Mb of chromosome 3 (File S2). We have chosen that region of chromosome 3 for not containing centromeres, 5S or 45S rRNA genes that due to natural variation in their copy number among accessions (Davison *et al.* 2007; Long *et al.* 2013) could affect our sequencing depth estimates.

Since the 134 individuals of the RIL population (Cvi-0 × Ler-0) were sequenced in separate Illumina lanes, we fitted a simple linear regression model on 18 technical replicates to account for the plate effect and obtain a single rRNA gene copy number estimate per line.

### Estimating rRNA gene copy number through qPCR

We estimated 45S rRNA gene copy number in the MA lines through quantitative PCR (qPCR) by comparing the abundance of the 18S rRNA subunit with the single copy gene At3g18780 (ACT2) according to:

rRNA gene copy number = 2^Ct(At3g18780 gene) − Ct(18S rRNA gene)^, where Ct(*x*) stands for the threshold cycle for *x*.

For the 18S rRNA gene we used primers 5'-CCT GCG GCT TAA TTT GAC TC-3' and 5'-GAC AAA TCG CTC CAC CAA CT-3', while for ACT2 primers 5'-TGC CAA TCT ACG AGG GTT TC-3' and 5'-TTA CAA TTT CCC GCT CTG CT-3' (Davison *et al.* 2007). We employed the FastStart Essential DNA Green Master kit (Roche) according to manufacturer's instructions in a LightCycler^®^ 96 (Roche) with the following thermal profile: preincubation at 95°C for 600 seconds; 45 cycles at 95°C for 10 seconds, 60°C for 15 seconds (in acquisition mode) and 72°C for 15 seconds; melting step at 95°C for 10 seconds, 65°C for 60 seconds and 97°C for 1 second. No primer dimers were detected in the melting curve.

4–5 biological replicates of MA lines 29, 39, 49, 59, 69, 79, 89, 99, 109 and 119 (Shaw *et al.* 2000; Ossowski *et al.* 2010) were propagated one generation by single-seed descent. We carried out qPCR of each line in 4 technical replicates (both for the 18S rRNA gene and ACT2) per plate. We distributed all lines in 14 96-well plates with some lines present in more than one plate. We included a common DNA control (accession id: 1002) to all plates for the purpose of standardization. Raw 18S rRNA gene copy number estimates and standardized values are provided in File S2. For the purpose of visualization we plotted 18S rRNA gene abundance relative to the lowest line mean value in generation 32 (line 69).

### Linkage mapping

Simple interval mapping (SIM) was performed with the R package R/qtl (Broman *et al.* 2003). Multiple QTL mapping (MQM) was done with a 2 centimorgan step size and 10 as window size (Arends *et al.* 2010). 1000 permutations were applied to estimate genome wide significance. QTL mapping in MAGIC lines and multiple imputation to determine estimated founder accession effects were performed with R/happy (Mott *et al.* 2000; Kover *et al.* 2009).

### CAPS analysis

Cleaved amplified polymorphic sequence (CAPS) analysis of RILs derived from the cross Cvi-0 x Ler-0 was performed as described elsewhere (Lewis *et al.* 2004). Briefly, DNA from each RIL was amplified by PCR in a 30 μl reaction with primers 5'-AGG GGG

GTG GGT GTT GAG GGA-3' and 5'-ATC TCG GTA TTT CGT GCG CAA GAC G-3', and the following thermal profile: 32 cycles at 95°C for 20 seconds, 62°C for 20 seconds and 72°C for 40 seconds. Resulting PCR products were incubated with restriction enzyme *Rsa*l (New England Biolabs Inc.) for 4hrs at 37°C and subjected to agarose gel electrophoresis. Cleaved PCR products correspond to Cvi-0 derived rRNA genes, while intact PCR products to Ler-0 derived rRNA genes. Results were summarized in File S2.

### Fluorescence in situ hybridization (FISH)

The preparation of root-tip meristem chromosome spreads followed the protocol published by Mandáková and Lysak (2016) (Mandáková and Lysak 2016). Seedlings were germinated on filter paper soaked in distilled water in a Petri dish at 21 °C. Cut, approx. 1 cm long, roots were pretreated with ice-cold water for ca. 24 hrs, then fixed in ethanol:acetic acid (3:1) fixative at 4 °C for 24 hrs. The fixed roots were rinsed in distilled water and 1x citrate buffer (10 mM sodium citrate, pH 4.8), and digested by 0.3% pectolytic enzymes (cellulase, cytohelicase and pectolyase) in 1x citrate buffer at 37 °C for 90 min. Individual root-tip meristematic tissues were dissected in ca. 20 μl of 60% acetic acid on a clean microscopic slide. Then the cell material was covered with a coverslip, evenly spread by tapping, and the slide gently heated over a flame. The slide was frozen in liquid nitrogen, coverslip flicked off, fixed in ethanol:acetic acid (3:1) fixative and air-dried. The suitable slides selected after inspection under a phase-contrast microscope were processed as described by Lysak and Mandáková (2013) (Lysak and Mandáková 2013). In brief, the slides were pretreated by ribonuclease A (100 μg/ml in distilled water) at 37 °C for 1 hr and by pepsin (0.1 mg/ml in 10 mM HCl) for at 37 °C for 1 – 3 min, and postfixed in 4% formaldehyde in 2x SSC (20x SSC: 3 M NaCl in 0.3 M sodium citrate, pH 7.0) at room temperature for 10 min. The slides were washed in 2x SSC between the steps and eventually dehydrated in an ethanol series (70%, 80%, and 96% ethanol, 3 min each).

*A. thaliana* BAC clone T15P10 containing 45S rRNA genes was used to identify the NORs. To identify *A. thaliana* chromosomes 2 and 4, eleven BAC clones from the upper arm of chromosome 2 (F2I9, T8O11, T23O15, F14H20, F5O4, T8K22, F3C11, F16J10, T3P4, T6P5, and T25N22) and 15 BACs from the upper arm of chromosome 4 (F6N15, F5I10, T18A10, F3D13, T15B16, T10M13, T14P8, T5J8, F4C21, F9H3, T27D20, T19B17, T26N6, T19J18, and T1J1) were used. The 45S rRNA gene probe was labeled with Cy3-dUTP, chromosome 2 BACs with biotin-dUTP and chromosome 4 BAC clones with digoxigenin-dUTP by nick translation (Lysak and Mandáková 2013). 100 ng from each labeled BAC DNA was pooled together, ethanol precipitated, dissolved in 20 μl of 50% formamide in 10% dextran sulfate in 2□ SSC and pipetted on the selected microscopic slides. The slides were heated to 80 °C for 2 min and incubated at 37 °C overnight. Hybridized DNA probes were visualised either as the direct fluorescence of Cy3-dUTP (yellow) or through fluorescently labeled antibodies against biotin-dUTP (red) and digoxigenin-dUTP (green). DNA labeling and fluorescence signal detection was carried out using a previously published protocol (Lysak and Mandáková 2013). Chromosomes and nuclei were counterstained with 4,6-diamidino-2-phenylindole (DAPI, 2 μg/ml) in Vectashield antifade. Fluorescence signals were analyzed and photographed using a Zeiss Axioimager epifluorescence microscope and a CoolCube camera (MetaSystems), and pseudocolored/inverted using Adobe Photoshop CS5 software (Adobe Systems). The size of fluorescence signals corresponding to the 45S rRNA gene probe was measured in Photoshop as a number of pixels per a defined area.

### Data Availability

DNA sequencing data from F2s and RIL populations have been deposited at the U.S. National Center for Biotechnology information(https://www.ncbi.nlm.nih.gov/bioproject) under BioProject: PRJNA326502. DNA sequencing data from the MAGIC lines and MA lines were downloaded from the European Nucleotide Archive (http://www.ebi.ac.uk/ena) under accession numbers PRJEB4501 and PRJEB5287 (Hagmann *et al.* 2015), respectively.

## Supplemental Material

**Figure S1.**
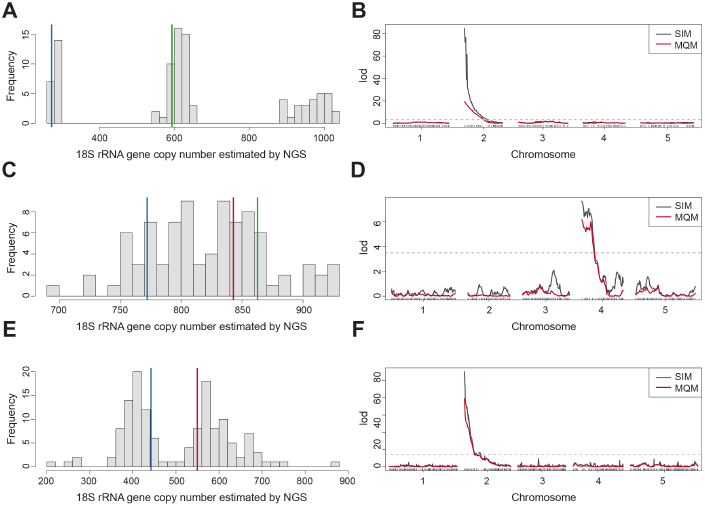
rRNA gene copy number variation in F2 progenies and a RIL population. (A) The distribution of 18S rRNA gene copy number estimated by NGS in an F2 population of 92 individuals derived from the cross Ull1-1 (8426) × TDr-7 (6193). Blue and green vertical lines represent phenotypic values of accession TDr-7 and an F1 individual, respectively (data for Ull1-1 are missing). (B) QTL mapping of 18S rRNA gene copy number in the same F2 population described in (A). (C) The distribution of 18S rRNA gene copy number estimated by NGS in an F2 population of 84 individuals derived from the cross T460 (6106) × Omn-5 (6071). Blue, green and red vertical lines represent phenotypic values of accession Omn-5, an F1 individual and accession T460, respectively. Note that the difference between the parental lines is small relative to the measurement error, and that this likely explains the “transgressive” value of the F1 individual. (D) QTL mapping of 18S rRNA gene copy number in the same F2 population described in (C). (E) The distribution of 18S rRNA gene copy number estimated by NGS in a RIL population of 134 individuals derived from cross Cvi-0 (6911) × Ler-0 (7213). Blue and red vertical lines represent phenotypic values of parental accessions Cvi-0 and Ler-0, respectively. (F) QTL mapping of rRNA gene copy number in the same RIL population described in (E).

**Figure S2.**
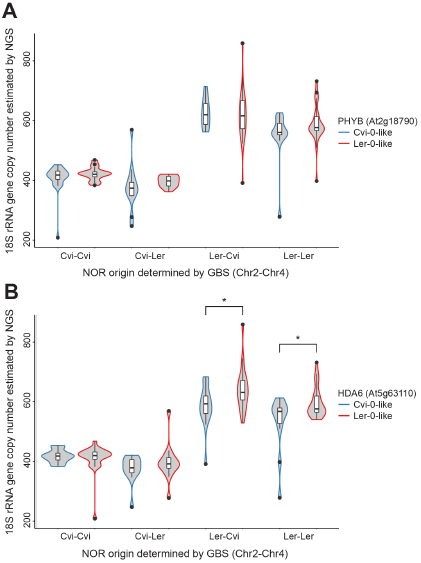
The effect of Cvi-0 and Ler-0 alleles at *PHYB* and *HDA6* genes in rRNA gene copy number. (A) 18S rRNA gene copy number in the Cvi-0 × Ler-0 RIL population estimated by NGS split — first — by NOR parental identity and — second — by the allele inherited at the *PHYB* locus as determined by GBS. Blue and red contours of the violin plots indicate alleles Cvi-0-like and Ler-0-like at the *PHYB* gene (At2g18790), respectively. P-values for the effect in rRNA gene copy number of parental NORs (NOR2-NOR4) Cvi-Cvi, Cvi-Ler, Ler-Cvi and Ler-Ler as a function of *PHYB* are 0.486, 0.897, 0.495 and 0.450, respectively. (B) 18S rRNA gene copy number in the Cvi-0 × Ler-0 RIL population estimated by NGS split — first — by NOR parental identity and — second — by the allele inherited at the *HDA6* locus as determined by GBS. Blue and red contours of the violin plots indicate alleles Cvi-0-like and Ler-0-like at the *HDA6* gene (At5g63110), respectively. P-values for the effect in rRNA gene copy number of parental NORs (NOR2-NOR4) Cvi-Cvi, Cvi-Ler, Ler-Cvi and Ler-Ler as a function of *PHYB* are 0.7038, 0.3731, 0.0298 and 0.0125, respectively. The asterisk (*) represent a p-value < 0.05.

**Figure S3.**
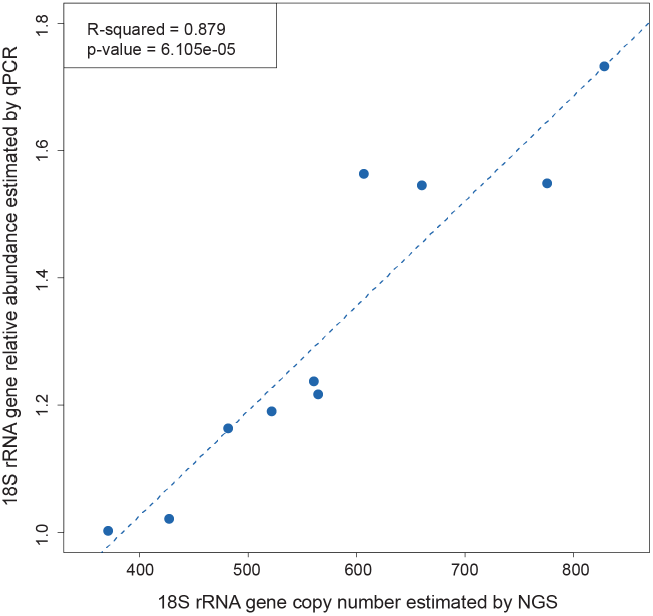
Correlation between two estimators of 18 rRNA gene copy number for the MA lines. Correlation between two estimators of 18 rRNA gene copy number for the MA lines: Next generation sequencing (NGS) and quantitative PCR (qPCR).

**Table S1.** Nucleolar association of NOR2 and NOR4 in two accessions. Relative frequency of root-tip nuclei with a particular NOR configuration relative to its close proximity to the nucleolus for the parental accessions Ale-Stenar-64-24 (1002) and TRÄ-01 (6244). n is the number of nuclei in each category.

| Accession name<br>(n = number of nuclei) | NORs associated with the nucleolus |  |  |  |  |
| --- | --- | --- | --- | --- | --- |
|  | NOR2 2x | NOR2 2x & | NOR2 2x & | NOR2 1x & | NOR4 2x |
|  |  | NOR4 1x | NOR4 2x | NOR4 2x |  |
|  | n (%) | n (%) | n (%) | n (%) | n (%) |
| Ale-Stenar-64-24 (n = 19) | 0 | 0 | 3 (16%) | 8 (42%) | 8 (42%) |
| TRÄ-01 (n = 25) | 10 (40%) | 14 (56%) | 1 (4%) | 0 | 0 |

**File S1. 45S rRNA gene reference.**

**File S2. Genetic maps of F2s and RILs, and phenotypes of F2s, RILs, MAGIC and MA lines.**

**File S3. Manuscript in preparation on NOR expression. F.A. Rabanal and M. Nordborg.**

## Acknowledgments

We sincerely thank Ashley Farlow and Ortrun Mittelsten Scheid for helpful discussions during the course of this project, Youssef Belkhadir for his valuable input in the preparation of this manuscript, and Claude Becker for providing seeds of the MA lines. This work was partly funded by a grant from the ERC (MAXMAP, Grant No. 268962).

## Author contributions

Conceived and designed the experiments: FAR, MN. Performed the experiments: FAR, VN, TM. Contributed reagents/materials/analysis tools: PN, MAL, RM. Wrote the manuscript: FAR, MN.

